# Long-term balancing selection maintains cryptic color polymorphism in frogs

**DOI:** 10.1101/2024.11.15.623899

**Authors:** Sandra Goutte, Stéphane Boissinot

## Abstract

Color polymorphism can influence the evolutionary fate of cryptic species because it increases populations’ chances of survival in heterogenous or variable environments. Yet, little is known about the molecular and evolutionary mechanisms underlying the persistence of cryptic color polymorphisms, or the impact these polymorphisms have on the macro-evolutionary dynamics of lineages. Here, we examine the evolutionary history of the most widespread cryptic color polymorphism in anurans, involving green and brown morphs. Using an order-scale comparative analysis, we show that these morphs can co-exist within species over long periods of evolutionary time and that polymorphic lineages switch habitat more frequently and have greater diversification rates than other groups. We then identify the locus responsible for the green/brown polymorphism in a group of African grass frogs, and demonstrate that this genomic region is evolving under long-term balancing selection, resulting in trans-specific polymorphism. These results provide a micro-evolutionary mechanism for the long-term persistence of color polymorphism observed at a macro-evolutionary scale. This study highlights the importance of cryptic color polymorphism in the ecology and evolution of anurans, and provides a framework for future research on the genetic architecture and selective forces underlying cryptic traits.

## Introduction

Animals exhibit a wide variety of colors and patterns that play crucial roles in their survival by helping them avoid detection by visual predators. These cryptic color traits allow individuals to blend into their surroundings, masquerade as inanimate objects like leaves or bird droppings, or disrupt their body shape^1^. The simultaneous occurrence of multiple color morphs within a species is common amongst cryptic-colored organisms, and similar color polymorphisms are found repeatedly across distantly-related species^2,3^. Cryptic color polymorphisms have inspired much theoretical and empirical work aimed at resolving the apparent paradox of phenotypic and genetic variation persisting despite genetic drift and various selective pressures tending to reduce or eliminate it^4–9^. Several evolutionary mechanisms such as frequency-dependent predation, variations in selection pressures across space or time, or heterozygote advantage, have been proposed^4,10–17^. Yet, it is often unclear whether selective processes can maintain multiple color morphs over long evolutionary periods or if color polymorphisms represent a transient state where one of the morphs is on its way to fixation^18^.

In anurans (frogs and toads), earthy tones that provide camouflage against vegetation or soil backgrounds are prevalent^19,20^. The most widespread form of color polymorphism in this group involves individuals displaying either a green, or a melanin-based, brown to grey, dorsal coloration — referred to hereafter as Green Color Polymorphism (GCP)^19,20^. Despite its commonness, the genetic basis and evolutionary mechanisms underlying anuran GCP remains poorly understood. To date, genetic investigations of anuran coloration have mostly focused on a handful of poison frog species, which display bright colors such as blue, red, or yellow, that serve aposematic and sexual signaling functions, and are rarely found in other taxa^21–23^. However, crossing experiments have revealed the mode of inheritance of colors in a broader taxonomic range^19^. For instance, crosses have established that GCP is controlled by a single locus with a dominant *Green* allele in three North American frogs^24–26^.

Here, we conduct an order-scale comparative analysis to retrace the evolution of GCP and show that this polymorphism can persist for long periods of evolutionary time and is associated with increased habitat transitions and diversification rates. We then investigate the genetic architecture of GCP in a radiation of African grass frogs endemic to the Ethiopian Highlands (12 species; genus *Ptychadena*). Within this radiation, four species (*P. robeensis*, *P. levenorum*, *P. erlangeri* and *P. nana*) form a monophyletic clade and share the same green/brown polymorphism^27^. Using whole-genome sequencing data, we identify and characterize the GCP locus and show that this genomic region is evolving under long-term balancing selection, resulting in trans-specific polymorphisms. We discuss how the genomic architecture of the trait and species’ ecology may play a role in the maintenance of cryptic color polymorphism, and how, in turn, GCP impacts the evolutionary fate of species carrying it.

## Results

### Green coloration evolves rarely and is more evolutionarily stable as a polymorphic state

To retrace the evolution of green coloration in anurans, we examined 13,025 photos of 2,363 species for which phylogenetic information was available^28^, representing 30.8% of species recognized at the time of writing^29^ (Supp. Table S1). We categorized species as “fixed green” (186 species), “polymorphic” (191 species), and “not green” (1,986 species), and fitted multiple k-states Markov models of evolution for green coloration (Supp. Table S2; Supp. Fig. S1). Models of evolution where transitions between the “fixed green”, “polymorphic”, and “not green” states occur at different rates best fitted our data. Additionally, the goodness of cit of the model allowing direct transitions between the “fixed green” and “not green” states (ARD model) was not significantly different from that of the one requiring an intermediate “polymorphic” state (POLY model; Supp. Table S2; Supp. Fig. S1). In both models, the rate of evolution of the green coloration was 20 times lower than the loss of fixed green coloration. The rate of fixation of green coloration was about half of that of its loss. In the full model where direct transitions between the “fixed green” and “not green” states were allowed, both transition rates were low, and the gain of green coloration (rate = 0.001) lower than its loss (rate = 0.008).

We estimated the number of changes between color states during the evolution of anurans based on 1,000 stochastic maps of the trait, allowing for all transitions to have different rates (Fig. 1A & 1B). We estimated that green coloration evolved 88 times (95% confidence interval: 69 – 104), was fixed 132 times (112 – 156) from a polymorphic state, and was lost 353 times (302 – 401). Direct transitions between fixed states had the lowest estimated number of occurrences, with 32 evolution (21 – 41) events and 27 losses (17 – 37). Across the phylogeny, the average evolutionary time spent in the “polymorphic” state was greater than in the “fixed green” state (Fig. 1C). These results reveal that GCP can persist over long periods of evolutionary time, which suggests the possible action of variation-maintaining selective forces.

**Figure 1.**
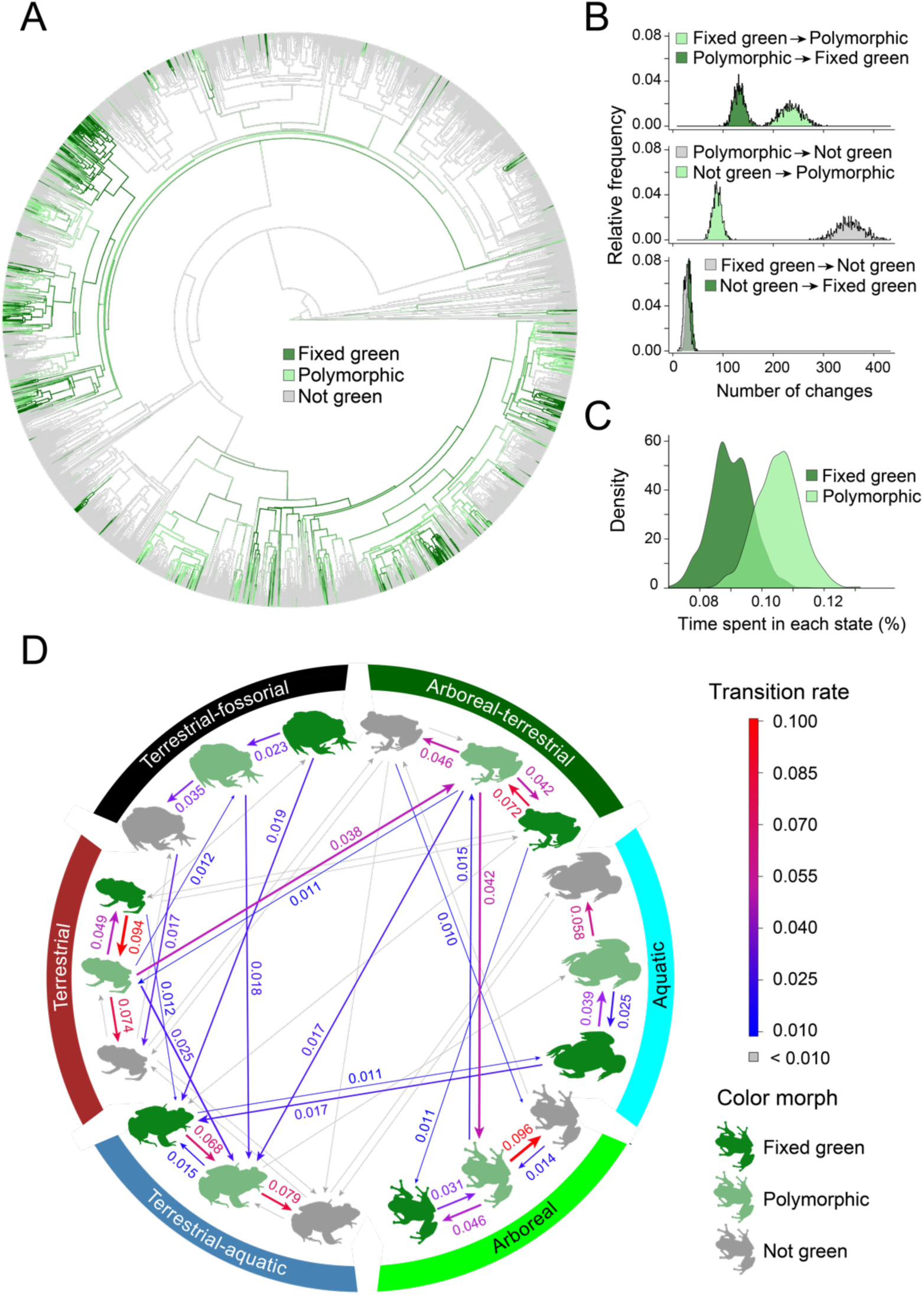
Evolution of the green coloration and GCP in anurans. **A** Ancestral state reconstruction of the green coloration and GCP in 2,363 anurans (30.8% of known anuran species, all families represented; phylogeny from^28^) based on 1,000 stochastic maps using a “all rates different” (ARD) model of evolution. **B** Number of transitions between color states (“fixed green”, “not green”, and “polymorphic”) during anurans’ evolutionary history, estimated based on 1,000 stochastic maps. **C** Estimated time spent in “fixed green” and “polymorphic” states during the evolutionary history of anurans, based on 1,000 stochastic maps. **D** Co-dependence of green coloration evolution and habitat transitions in anurans. Transition rates estimated by the best fitting model of joint color and habitat evolution (see text) and above 0.010 are given on the arrows.

### Color polymorphism is associated with higher habitat transition rates

To determine whether the evolution of GCP is driven by environmental factors, we assembled habitat data for 1,989 species and compared the cit from multiple models of joint evolution between coloration and habitat states. A habitat-dependent model for the evolution of green coloration provided the best cit for our data (Supp. Table S3; Fig. 1D). As in our global model of evolution, the rate of loss of green coloration was greater than its evolution rate, in all habitats. However, in arboreal, arboreal-terrestrial, and terrestrial lineages, the fixation rate of green coloration was substantial (> 0.040). Interestingly, transition rates between habitats were higher in “polymorphic” lineages compared to other groups (Fig. 1D). In particular, transitions rates were the highest in GCP lineages that transitioned from terrestrial to terrestrial-aquatic (0.025) or arboreal-terrestrial (0.038), and from arboreal-terrestrial to arboreal (0.042). These results indicate that green coloration is differentially selected in different habitats, and that GCP may facilitate habitat transitions.

### Diversification rate is higher in polymorphic lineages

Theoretically, color polymorphism has the potential to facilitate speciation and reduce extinction risk^8,18,30^, but few empirical studies have assessed the relationship between color polymorphism and diversification rate thus far^31–34^. To test whether GCP is associated with increased speciation and reduced extinction in anurans, we compared GCP- dependent and independent models of diversification on the 5,242 species included in the largest molecular anuran phylogeny available, representing 68.3% of known taxa^28^. A GCP-dependent model best fitted our data (Supp. Table S4). In this model, speciation rate (λ) was greater and extinction rate (µ) lower in “polymorphic” lineages compared to “fixed green” and “not green” groups (Table 1). Extinction rate was the highest in “fixed green” lineages, while their speciation rate was roughly half of that of “polymorphic” lineages, but twice of that of “not green” groups. These results indicate that GCP is associated with an increased net diversification rate in anurans.

**Table 1.** Coefficients for the full speciation/extinction (MuSSE) model. The FULL MuSSE model better fitted our data compared to color state-independent (NULL), equal rates (ER MuSSE) and color state-independent and equal rates (ER NULL) models (see text and Supp. Table S4).

| | Speciation rate<br>($\lambda$) | Extinction rate<br>($\mu$) | Transition rate<br>($q$) |
| --- | --- | --- | --- |
| Fixed green | 0.1171 | $1.89 \cdot 10^{-6}$ | |
| Polymorphic | 0.2393 | $3.42 \cdot 10^{-8}$ | |
| Not green | 0.0558 | $9.18 \cdot 10^{-7}$ | |
| Fixed green →<br>Polymorphic |  |  | 0.0743 |
| Polymorphic → Fixed<br>green |  |  | 0.0136 |
| Polymorphic → Not green |  |  | 0.2565 |
| Not green → Polymorphic |  |  | 0.0084 |

### A single genomic region is associated with GCP in the African grass frog Ptychadena robeensis

Our macroevolutionary analysis indicates that GCP may persist for long periods of evolutionary time, suggesting that selection may maintain this polymorphism. In order to determine a possible role of selection, we focused on the African grass frog *Ptychadena robeensis*, where individuals within the same population exhibit either a green or a brown dorsal coloration (Fig. 2A). In this species, green and brown skins differ in their density of iridophores, a cell type containing crystals that reflect white-to-blue colors (Supp. Methods; Supp. Fig. S2). Additionally, guanine crystals contained within the iridophores are larger and rounder in green skin compared to brown skin, which may explain their difference in reflectance^35^ (Supp. Fig. S2).

**Figure 2.**
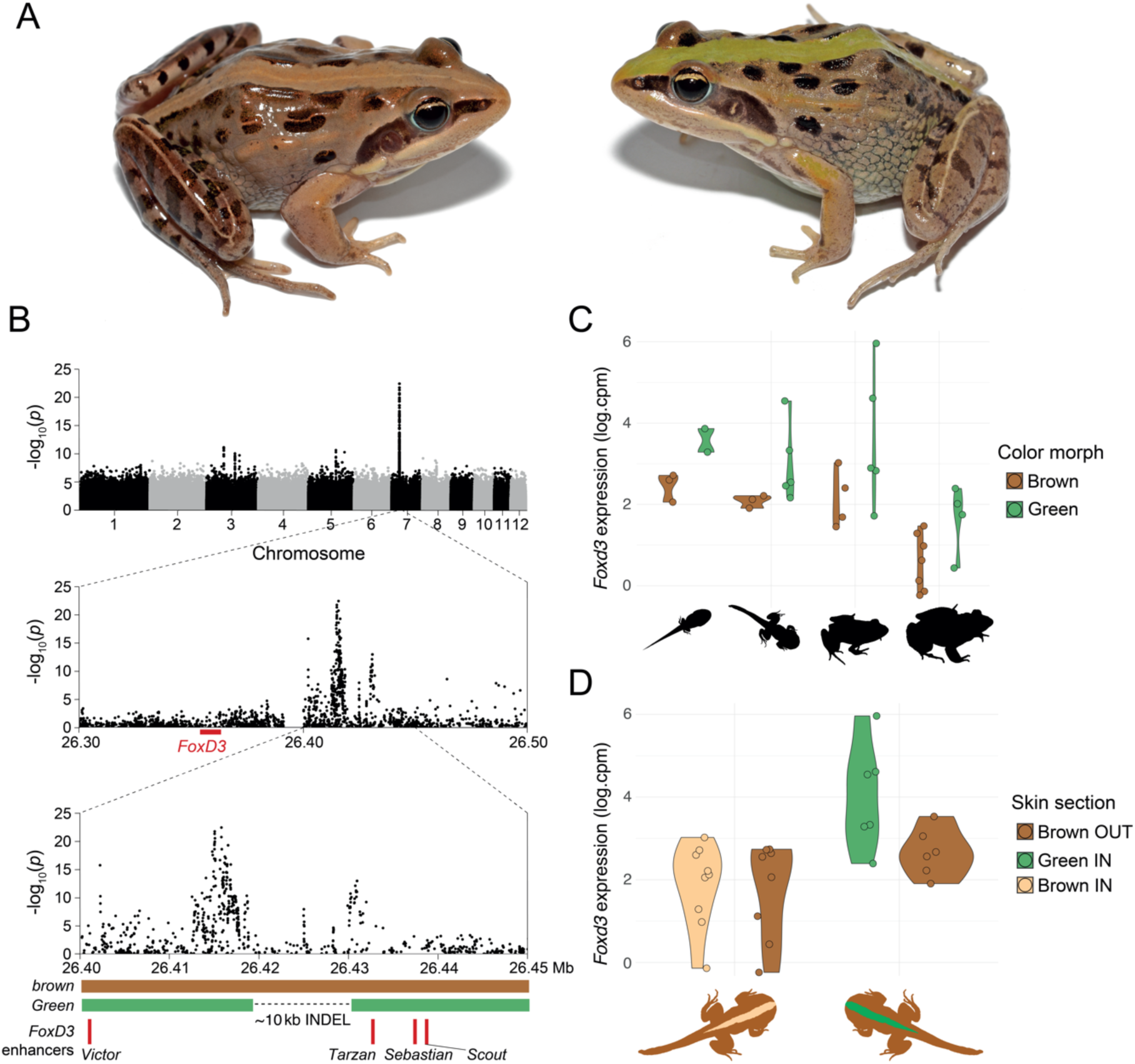
A single locus near *FoxD3* is associated with dorsal coloration in *Ptychadena robeensis*. **A** Brown (left) and green (right) morphs in *Ptychadena robeensis*. **B** Genome-wide association study of dorsal coloration in *Ptychadena robeensis* contrasting brown (n=61) with green individuals (n=39). The *Green* and *brown* alleles, as well as the known *FoxD3* enhancers in the region are represented below the Manhattan plots. **C** Expression levels of *FoxD3* in dorsal skin of *Ptychadena robeensis* (n=33) with green and brown genotypes at different development stages. From left to right: tadpole (3 brown, 2 green), metamorph (3 brown, 5 green), juvenile (4 brown, 5 green), and adult (7 brown, 4 green). **D** Expression levels of *FoxD3* in dorsal skin of *Ptychadena robeensis* (n=28) in wide-striped, green and brown genotyped frogs within (8 brown, 6 green) and outside (8 brown, 6 green) the stripe.

We conducted a genome-wide association study (GWAS) to identify the genetic basis of GCP in *Ptychadena robeensis*, for which a chromosome-level reference genome is available^36^. We obtained whole genome resequencing data from 100 *P. robeensis* (average 2.78X coverage), including 61 brown and 39 green individuals. We found a single association peak on chromosome 7, located 52 kb upstream of the gene *FoxD3* between two known *FoxD3* enhancers, *Tarzan* and *Victor*^37^ (Fig. 2B). Examination of this genomic region across the population established that the *Green* allele is dominant over the *brown* allele, and that most green individuals are heterozygous at this GCP-associated locus. Additionally, the *Green* haplotype carries a ∼10 kb deletion adjacent to the association peak (Fig. 2B). The gene *FoxD3* is known to interact with the transcription factor *Mitf*^38^ and is involved in iridophore differentiation in zebrafish, where an increase in *FoxD3* expression leads to differentiation of chromatophore precursors into iridophores instead of melanophores^39^. Thus, given the location of our association peak near known *FoxD3* enhancers, we hypothesized an increased expression of *FoxD3* in the skin of green frogs.

### FoxD3 is differentially expressed in green versus brown skin

We compared *FoxD3* expression levels in the dorsal skin of 33 *P. robeensis* individuals across four developmental stages: tadpole, metamorph, juvenile and adult. For tadpoles and metamorphs, where adult coloration has not developed, we examined whole-genome resequencing data and used their genotype at the GCP locus to predict future coloration. *FoxD3* expression was higher in green *versus* brown-genotyped individuals, across all developmental stages (t = -2.43, p-value = 0.02), and was overall higher in developing frogs compared to adults (Fig. 2C & Supp. Table S5). Additionally, we compared *FoxD3* expression levels between skin patches in the vertebral stripe that were or would become green, with skin patches of the same individuals outside of the stripe that were and would remain brown (n = 14; Fig. 2D). We found that *FoxD3* expression was elevated in the skin that was or would become green but not in the areas that would remain brown (Supp. Table S5). These results indicate that an increase in *FoxD3* expression level is associated with the appearance of green coloration *in P. robeensis*, and that expression of *FoxD3* in skin cells is both temporally and spatially modulated within individuals.

### The Green and brown alleles are maintained by balancing selection in P. robeensis

The simultaneous occurrence of two alleles within a species could be the result of incomplete fixation of one of the alleles or could be maintained by balancing selection through heterozygote advantage, negative frequency-dependent selection, or heterogeneity in selection pressure across time or space^40^. Balancing selection, if ancient enough, leaves genomic signatures distinguishable from other evolutionary processes^41^. We searched for signatures of selection in the genome of *Ptychadena robeensis* using data from 10 frogs sequenced at higher-coverage (8.33X). We found a peak of genetic diversity (π) corresponding to our association peak in the green individuals (mostly heterozygotes), while sequences in that region were all nearly identical in the brown frogs (Fig. 3A). The association peak also coincided with a peak in absolute sequence divergence (D_XY_) and genetic differentiation (F_ST_) between the two morphs (Fig. 3B & 3C), suggesting an ancient divergence of the two alleles. We then computed two statistics specifically designed to test for balancing selection, Tajima’s D, which compares the number of segregating sites and nucleotide diversity in a genomic region^31^, and β, which is based on correlations of allele frequencies across neighboring loci^32^. Tajima’s D was positive across the GCP locus, indicating an increase in intermediate frequency polymorphism, which is consistent with balancing selection (Fig. 3D). We also found a peak of β scores (above the 99^th^ genome-wide percentile) matching our association peak (Fig. 3E), which strongly suggests that long-term balancing selection is acting on the GCP locus and maintains the *Green* and *brown* alleles in *Ptychadena robeensis*.

**Figure 3.**
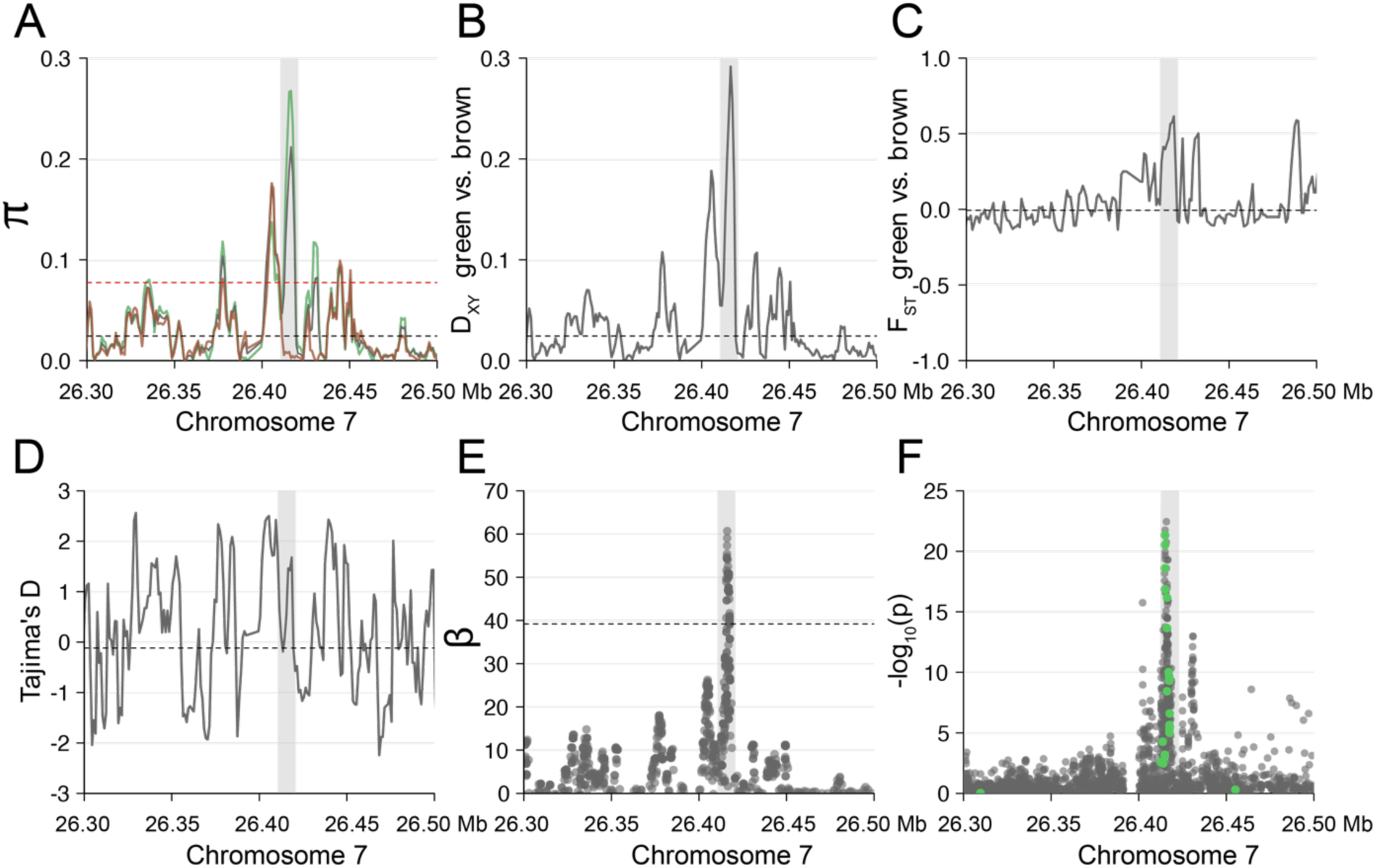
Signatures of balancing selection in the GCP region of *Ptychadena robeensis*. All summary statistics (**A-E**) were computed on 3 kb sliding windows with 1 kb overlap on the higher-coverage dataset (n=10, 8.33X average coverage). Grey-shaded areas correspond to the GCP-associated region (chromosome 7, position: 26,411,500- 26,422,300). Black horizontal dashed lines represent the genome-wide average statistics’ value. **A** Population-wide (grey), brown individuals (brown; all are homozygotes), and green individuals (green; most are heterozygotes) nucleotide diversity (π). Red horizontal dashed line represents the genome-wide 95^th^ percentile of nucleotide diversity. **B** Absolute genetic divergence (D_XY_) between green and brown individuals. **C** Relative differentiation (F_ST_) between green and brown individuals. **D** Tajima’s D values across the GCP region. **E** β scores. Red horizontal dashed line represents the genome-wide 99% percentile of β scores. **F** Genome-wide association study (GWAS) contrasting brown (n=61) and green (n=39) individuals. Green full circles represent individual SNPs which are polymorphic in all four species of the *Ptychadena erlangeri* group (*P. robeensis, P. nana, P. levenorum*, and *P. erlangeri*).

Local suppression of recombination can be instrumental for the persistence of balanced polymorphisms by preserving the integrality of divergent haplotypes generation after generation^40^. This phenomenon is observed, for example, in alleles originating in chromosomal inversions^3,40,42^. Surprisingly, in *Ptychadena robeensis*, we found an elevated recombination rate at the GCP locus compared to the chromosome-wide average (Supp. Fig. S3). This pattern could reflect the long divergence time between the alleles, during which nucleotide diversity may have accumulated in the region surrounding the selected locus^43^.

### Long-term balancing selection maintains trans-species color polymorphism in the Ethiopian Highlands grass frog radiation

To establish how long balancing selection has maintained color polymorphism in the Ethiopian Highlands grass frog radiation, we investigated whether the alleles found in *P. robeensis* were shared with the other three polymorphic *Ptychadena* species of the clade (*P. nana, P. levenorum,* and *P. erlangeri*). We found that variation at the same locus was associated with coloration in all three species, with a large number of shared nucleotide polymorphisms (Fig. 3F). Additionally, we found elevated nucleotide diversity (π) and sequence divergence (D_XY_) between green and brown individuals at the GCP locus in all three species, as well as elevated F_ST_ between green and brown individuals for *P. nana* and *P. erlangeri* (Supp. Fig. S4). We reconstructed a maximum-likelihood haplotype phylogeny of GCP locus (3.38 kb; see Methods) including all 12 species of the Ethiopian Highlands *Ptychadena* radiation (Fig. 4). The haplotype phylogeny shows a well-supported divergence of the *Green* and *brown* haplotypes, preceding the divergence of the four polymorphic *Ptychadena* species, estimated ∼8 million year ago^44^ (Fig. 4). According to our phylogenetic reconstruction, the *Green* and *brown* haplotypes divergence even precedes the split between the *P. erlangeri* species group and the monomorphic (brown) *P. neumanni* species group, estimated ∼9-10 million year ago^44^ (Fig. 4). This result represents a strong evidence for ultra long-term balancing selection (as defined in^41^) maintaining the *Green* and *brown* alleles in these four *Ptychadena* species.

**Figure 4.**
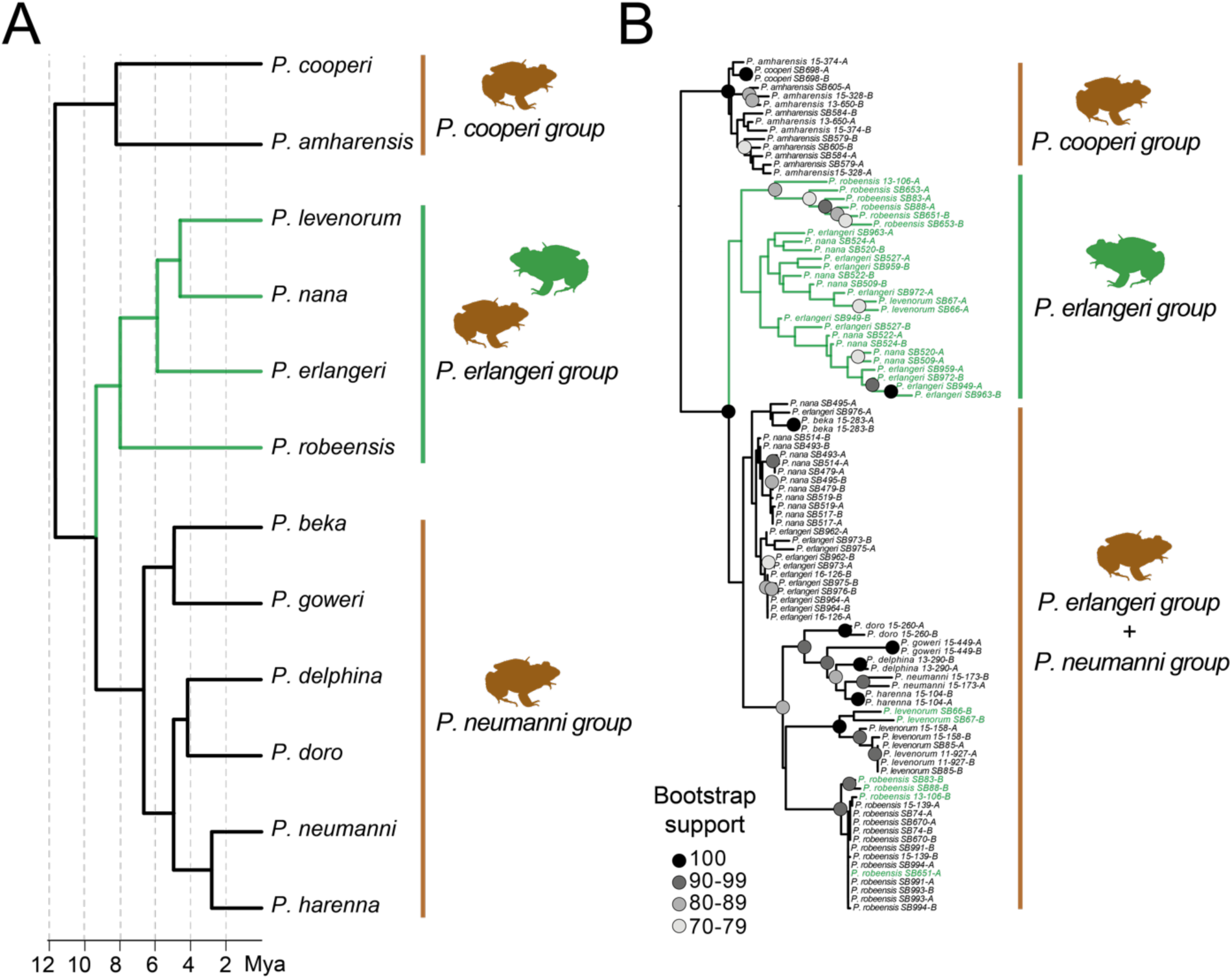
The *Green* and *brown* alleles are under long-term balancing selection in four species of Ethiopian *Ptychadena*. **A** Species tree including all 12 species of the Ethiopian Highlands *Ptychadena* radiation with estimated divergence times, adapted from^44^. **B** Raxml-ng tree with 100 bootstrap iterations of the region of interest (3,383 bp; 762 SNPs after filtering) on the phased haplotypes of the 12 *Ptychadena* species. Haplotypes of individuals with a green phenotype are indicated in green. One of the haplotypes of four green *P. robeensis* and two green *P. levenorum* are grouped in the “brown” clade, indicating that those individuals are heterozygous at the GCP locus.

## Discussion

Widespread GCP in anurans allowed us to investigate the mechanisms maintaining cryptic color polymorphism at multiple scales, and evaluate its evolutionary consequences. Within the Ethiopian Highlands *Ptychadena* radiation, we showed that the green and brown color morphs have been maintained for millions of years by long-term balancing selection, leading to trans-specific polymorphism. At the scale of the order, we found that GCP can persist for long periods of evolutionary time and is associated with frequent habitat transitions and high net diversification rates. Our results suggest that long-term balancing selection maintaining cryptic color polymorphism might be more prevalent than previously thought, and that GCP might play an important role in the eco-evolutionary fate of anuran species carrying it. Below, we discuss three aspects of our results: (1) how genomic architecture of cryptic color traits may favor the maintenance of polymorphism through balancing selection, (2) the possible ecological drivers of this balancing selection, and (3) the macroevolutionary consequences of cryptic color polymorphisms maintenance in anurans.

### The genetic architecture of GCP favors the maintenance of polymorphism

How the genomic architecture of a trait influences its evolutionary fate and how, selection, in turn, shapes the trait’s genetic makeup is a fundamental question in evolutionary biology. If a polymorphic trait is governed by multiple, unlinked loci, high gene flow within a population is likely to produce intermediate, sub-optimal morphs through recombination^40,45^. Single-locus architectures, such as the one governing GCP in the Ethiopian Highlands *Ptychadena* and other anurans^19^, thus favor the maintenance of polymorphism. Dominance also impacts the probability of polymorphism maintenance. Because it is dominant over the ancestral *brown* allele, the newly-evolved *Green* allele is likely to invade a population rapidly if the green phenotype offers fitness benefits, particularly under negative frequency-dependent selection regimes, where rare phenotypes are selected for^46,47^. After the initial invasion, the dominance of *Green* over *brown* in turn favors the maintenance of polymorphism within populations by maintaining the recessive *brown* allele in heterozygotes, even when the green phenotype is selected for. The single-locus, dominant/recessive biallelic system determining dorsal coloration in the Ethiopian Highlands *Ptychadena* thus represents a favorable genomic architecture for the maintenance of polymorphism.

### Possible ecological drivers of balancing selection in the Ethiopian Highlands grass frogs

Even with a favorable genomic architecture, long-term maintenance of polymorphism is considered unlikely because directional selection and drift are expected to lead to the fixation or loss of any allele over time^48^. The maintenance of the *Green* and *brown* alleles over millions of years of divergence and multiple speciation events in *Ptychadena* thus represents strong evidence for long-term balancing selection. Differential selective values of alternative morphs across space and/or time constitutes the simplest explanation for balanced cryptic color polymorphism in natural populations^6,49,50^. As selection by visual predators is expected to impose a strong pressure on anuran cryptic color evolution^19^, predators’ cognition and the level of crypsis in a given habitat should predict color morphs’ adaptive values at any point in time and space, and may explain the simultaneous presence of multiple morphs within a population. Individuals of all four polymorphic Ethiopian Highlands *Ptychadena* inhabit heterogenous habitats consisting of grasslands and cultivated fields, and are found on brown (soil, mud) or green (grass) substrates, where the green and brown morphs offer different levels of crypsis. Predators of these species likely include birds and mammals, which can form searching images and may favor commonly-encountered morphs, leading to apostatic selection^6^. Both ecological heterogeneity and negative frequency-dependent selection by visual predators promote cryptic color polymorphism in prey^49^, and may thus explain the maintenance of the green and brown morphs in the *Ptychadena erlangeri* species group, although experimental data would be needed to confirm this hypothesis.

Balancing selection may also act upon other trait(s) correlated with GCP by pleiotropy. Traits that have been found to be associated with color morphs in anurans include female fecundity^51^, overwinter survival^52^, immunity^10^, and larval development rate^53^. In these systems, selection can act in opposite directions on the color trait and a pleiotropic trait, or two color-associated traits. For example, in the cricket frog *Acris crepitans,* individuals with the advantageous green-or red-striped morphs have lower immune capabilities compared to grey individuals, which may lead to balancing selection of the morphs^10^. Polymorphism can also be maintained if heterozygotes have a fitness advantage compared to either homozygote. Heterozygote advantage can arise in genomic regions with low recombination and/or in which the alleles are old, because deleterious mutations may have accumulated on haplotypes underlying rare morphs, in particular if the haplotype is dominant and mostly found in heterozygotes^40^. In the Ethiopian Highlands grass frogs, color polymorphism is associated with variation in the regulatory region of *FoxD3*, a gene not only involved in chromatophore differentiation^39^, but also in neural crest development and autoimmune diseases in humans^54^. It is therefore possible that the *Green* allele, mostly found in heterozygotes, accumulated recessive mutations impacting other functions of *FoxD3* that are deleterious in homozygote *Green* individuals.

### Ecological and evolutionary implications of cryptic polymorphism maintenance in anurans

In our anuran macroevolution analysis, we found that GCP is associated with increased habitat transition rates, supporting the hypothesis that cryptic color polymorphism allows greater adaptability to novel ecological niches and therefore promotes range expansion^8,55^. In agreement with classical ecological evolutionary theory^8,18,30,56^, we found a higher diversification rate in GCP lineages compared to “fixed green” and “not green” groups. This result suggests that colonization of new environments and biogeographic areas by polymorphic populations leads to greater speciation, and lower extinction rates. Several mechanisms can explain the relationship between GCP, habitat transition, and diversification. Speciation rate may increase when polymorphic populations become monomorphic in the process of colonizing new ecological niches, resulting in rapid divergence and reproductive isolation from ancestral populations^31^. Cryptic color polymorphisms may also promote the persistence of populations under variable environmental conditions, which in turns lowers their risk of extinction^8,55^. Finally, rapid speciation may favor color polymorphism persistence by providing opportunities for the reintroduction of lost ancestral morphs through adaptive introgression, or their maintenance through the persistence of the ancestral selection regime in recently-evolved daughter species^45^.

To our knowledge, the association between color polymorphism and diversification rate at a macroevolutionary scale has only been examined in non-passerine birds^31^, Lacertidae lizards^32,33^, and carnivores^34^ so far. These studies yield mixed results: in carnivores and three of the five bird families studied, accelerated speciation rates were linked to color polymorphism ^31,34^, while in lacertids, two studies using the same species but different morph coding methods produced opposite results^32,33^. Interestingly, in birds of prey, where color polymorphism is associated with higher speciation rates, polymorphic species tend to have larger geographic ranges and occupy more diverse environments than monomorphic species. The clear patterns we obtained in our macroevolution and diversification analyses including over 2,300 species across the order suggest that consistent forces drive the evolution and maintenance of GCP in anurans. These results provide a foundation for future research on how color polymorphism may facilitate dispersal to new ecological niches and enhance the survival of populations facing environmental changes.

## Acknowledgements

We thank EWCA and EBI for assistance with sampling and export permits. We thank all the students, scientists and local collaborators who have contributed to sample collection over the years, and, in particular, Abeje Kassie, Megersa Kelbessa, Itbarek Kibret, and Samuel Woldeyes. We are grateful to Marc Arnoux, Nizar Drou, Rachid Rezgui, Sneha Thomas, Rainer Straubinger, David Howse and Sayel Daoud for lab assistance. Andrew Shantz, Mariana Lyra, and Theo Busschau provided helpful comments on the manuscript. This project was funded by NYUAD Grant AD180 to S.B. Sequencing and microscopy were carried out using the Core Technology Platforms resources at New York University Abu Dhabi. Analyses were carried out using the High-Performance Computing resources at NYUAD. Bioinformatics assistance was provided by the NYUAD Bioinformatics Core at New York University Abu Dhabi, supported by the Center for Genomics and Systems Biology. This work was partially supported by Tamkeen under the NYU Abu Dhabi Research Institute Award to the NYUAD Center for Genomics and Systems Biology (ADHPG-CGSB).

## Material and Methods

### Order-scale comparative analysis

We conducted a comparative analysis using the most recent molecular phylogeny for anurans, which comprises 5,242 species, representing all families and 68.5% of species recognized at the time of writing^28,29^. We collected data on dorsal coloration for 2,363 species, examining all photographs available for each species on Amphibiaweb (https://amphibiaweb.org). When no or few photos were available, we searched additional sources such as original species descriptions and the number of photographs examined for each species was systematically recorded (Supp. Table S1). In total, we examined 13,025 photos. Dorsal coloration was categorized as either “fixed green” (green coloration always present), “not green” (green coloration always absent), or “polymorphic” (green coloration present in some individuals only). Frogs with translucent green coloration, such as the glass frog *Hyalinobatrachium vireovittatum* for example, were not coded as “fixed green” or “polymorphic”, since the color is known to be of different molecular origin and likely plays a different ecological role^57^.

Habitat use data was collected independently for 1,989 species based on multiple large studies, and species accounts in Amphibiaweb and the IUCN red list websites (https://amphibiaweb.org; https://www.iucnredlist.org; Supp. Table S1). We categorized anuran habitats as arboreal, aquatic, terrestrial, arboreal/terrestrial, terrestrial/fossorial, and terrestrial/aquatic, based on the main habitat occupied by adult individuals (when reproductive and general habitats were available). We searched for precise microhabitat use descriptions such as “Individuals are found on roots, leaf litter and crevices” or “Adults are found in phytotelmas up to four meters height”, and considered insufficient general habitat description such as “This species inhabits riparian forests” or “This frog is linked to streams” as they do not provide sufficient details on whether adults spend their time on vegetation, on the ground or in the water.

### Evolution of the green coloration in anurans

We fitted multiple k-stats Markov (Mk) models of evolution for the green coloration on our 2,363 species dataset and the accordingly pruned phylogeny from^28^ using the function *;itMk* in R package *phytools*^58^. We compared four models of evolution: equal transition rates among “fixed green”, “not green” and “polymorphic” states (ER; one parameter), all transition rates different (ARD; six parameters), symmetric transition rates (SYM; three parameters), and a model allowing only transitions between green and not green through the “polymorphic” state (POLY; four parameters). We compared the cit of our models by computing AIC and Akaike weights, and conducting pairwise likelihood ratio tests. The ARD and POLY models better fitted the data compared to the ER and SYM models. Goodness of cit was marginally greater for the ARD compared to the POLY model, but it was not significantly different when accounting for the number of parameters included for each model. We then reconstructed color ancestral states by creating 1,000 stochastic maps of the trait onto the phylogeny using the ARD model and the function *make.simmap* to estimate the number of transitions between states and the time spent in each state across the phylogeny.

### Habitat-dependent evolution of the green coloration and GCP in anurans

To assess whether the evolution of green coloration and GCP were habitat-dependent, we first evaluated which model of evolution for habitat preference best fitted our data (Supp. Fig. S5). We compared the goodness of cit of Mk models with all transition rates between habitats equal (ER; 1 parameter), all transition rates different (ARD; 30 parameters), transition rates different for each habitat pair but equal in either direction (SYM; 15 parameters). The ARD model better fitted our data compared to the ER and SYM models. In an attempt to reduce the number of estimated parameters, we compared three additional models representing reduced versions of the ARD model: the ordered model, allowing transitions between adjacent habitats (e.g. the transition fossorial-terrestrial -> arboreal was not allowed in this model). This model allowed the following transitions: arboreal ˂-˃ arboreal-terrestrial ˂-˃ terrestrial ˂-˃ terrestrial-aquatic ˂-˃ aquatic and terrestrial ˂-˃ fossorial-terrestrial (Supp. Fig. S5). The two other models considered only transition that were estimated ≥ 0.001 (Reduced model 1), and > 0.001 (Reduced model 2), in the ARD model. All models were cit on the 1,989 species dataset using the *;itMk* function. The reduced model 2 fitted our data substantially better than all other models (Supp. Table S6; Supp. Fig. S6), we thus used this model in subsequent analyses.

To test for co-evolution between color states and habitats, we created a variable combining both variables for each species (e.g., “fixed green”+ arboreal, “polymorphic”+ terrestrial, etc.). We then fitted multiple models of evolution using the *;itMk* function. To limit the number of estimated parameters, we only allowed transitions permitted in our best-fitting model for habitat evolution (reduced model 2), from “fixed green” to “not green” and vice-versa through the “polymorphic” state, and did not allow simultaneous habitat and color states transitions (Supp. Fig. S7). The most complete model allowed all transition rates to be different (FULL model; 60 parameters). Model 2 (16 parameters) allowed different transition rates between habitats and between color states, but habitat transitions rates were equal across color states and color state transitions were independent of habitat. Model 3 (38 parameters) allowed all color state transition rates to be different within and across habitats, but transitions between habitats had the same rate regardless of the color state. Model 4 (40 parameters) allowed all color state transition rates to be different, but independent of habitat, and habitat transitions rates to be all different and vary according to color state (Supp. Fig. S7). We compared models’ cit by computing their respective AIC and Akaike weights. The FULL model best fitted our data (Supp. Table S3).

### Color state-dependent diversi;ication analysis

To detect potential association between color state (“fixed green”, “not green” or “polymorphic”) and speciation and/or extinction rates, we compared the goodness of cit of color state-dependent and color state-independent diversification models using the *make.musse* function implemented in the R package *diversitree*^59^. For this analysis, we used the complete tree from^28^ comprising 5,242 anuran species and our full dataset of 2,363 characterized species. We fitted four models: a model where speciation and extinction rates are independent of color state, and in which transition from “fixed green” to “not green” and vice-versa have to go through the “polymorphic” state (NULL; 6 parameters); a further simplified model in which transition rates between color states are all equal (ER NULL; 3 parameters); a model in which speciation and extinction rates are color state-dependent but transition rates between color states are all equal (ER MuSSE; 7 parameters); a model with color state-dependent speciation and extinction rates and all transition rates different (MuSSE; 10 parameters). We compared the goodness of cit of our models by computing their respective AIC and Akaike weights and conducting a likelihood ratio test against the simplest model (ER NULL; Supp. Table S4).

### Sampling of Ethiopian Ptychadena

Individuals of the Ethiopian Highlands *Ptychadena* radiation were collected between 2011 and 2022. Our study was approved by the relevant Institutional Animal Care and Use Committee at Queens College and New York University School of Medicine (IACUC; Animal Welfare Assurance Number A32721–01 and laboratory animal protocol 19–0003). Frogs were sampled according to permits DA31/305/05, DA5/442/13, DA31/454/07, DA31/192/2010, DA31/230/2010, DA31/7/2011, and DA31/02/11 provided by the Ethiopian Wildlife Conservation Authority. We photographed individuals in life and euthanized them by ventral application of 20% benzocaine gel. We extracted tissue samples and stored them in RNAlater or 95% ethanol. Adult individuals were fixed in 10% formalin for 24–48 h, and then transferred to 70% ethanol. After preservation, we took additional photographs of all individuals. All specimens were deposited at the Natural History Collection of the University of Addis Ababa, Ethiopia. Tissue samples are deposited at the Vertebrate Tissue Collection, New York University Abu Dhabi (NYUAD).

### Ptychadena robeensis reference genome

The assembly and annotation of the *Ptychadena robeensis* genome is described in details elsewhere (Hariyani et al. *in prep.).* Briefly, a *de novo* assembly was first constructed using two paired end libraries of mean insert size ∼350 bp and one paired end library of ∼550 bp insert size. The shotgun reads were incorporated with low-coverage PacBio to generate a hybrid assembly. To achieve longer scaffolds, two Chicago libraries were sequenced as described in^60^. The input hybrid assembly and Chicago library reads were then used as input data for HiRise, a software pipeline designed specifically for using proximity ligation data to scaffold genome assembly^60^. Finally, an Omni-C library was sequenced. The Omni-C reads and the input de novo assembly from the previous step were used as input data for HiRise. The resulting genome is 1.59 Gb in length and consists of 12 scaffolds, which corresponds to the haploid chromosome number reported in the genus *Ptychadena*, with an N50 of 157 Mb.

### DNA and RNA Extractions and Sequencing

Genomic DNA of 120 *Ptychadena robeensis,* 10 *P. nana,* 5 *P. levenorum,* 11 *P. erlangeri,* 6 *P. amharensis* individuals, as well as of one individual of *P. beka, P. cooperi, P. delphina, P. doro, P. goweri, P. harenna,* and *P. neumanni,* was extracted from liver or muscle tissue using the DNeasy blood and tissue kit (Qiagen, Valencia, CA; Supp. Table S7 & S8). RNA was extracted from the skin of 33 individuals using a RNeasy mini kit (Qiagen, Valencia, CA). For 14 wide-striped individuals, RNA was extracted from dorsal skin within and outside the vertebral stripe separately. For all other individuals, RNA was extracted from dorsal skin (Supp. Table S9). We quantified extracted DNA and RNA using a Qubit fluorometer (Life Technologies). Libraries were prepared using a NEB library prep kit and sequenced on Illumina NextSeq 550 flow cells at the Genome Core Facility of New York University Abu Dhabi. After quality filtering, reads were aligned to the *Ptychadena robeensis* reference genome (Hariyani et al. *in prep*.). The average coverage of the *P. robeensis* genomic data was 2.78X, except for ten individuals which we sequenced at an average of 8.33X. Other *Ptychadena* species were sequenced at an average of 6.80X, except for *P. delphina*, for which coverage was lower (3.50X). Variants were called using the function *HaplotypeCaller* from *gatk v3.5*^61^. The low-coverage *P. robeensis* and higher-coverage *P. robeensis* together with the other *Ptychadena* samples were then combined and genotyped in two separate datasets using *CombineGVCF* and *GenotypeGVCFs* functions from *gatk*.

### Genome-wide Association Study in Ptychadena robeensis

We examined the low-coverage genomic dataset (*n* = 106 individuals; SNPs PCA, admixture analysis, and visual examination), and removed hybrids resulting from the crossing of *P. robeensis* and *P. levenorum*, a closely related species with a partially overlapping distribution range^2,27^. We then used VCFtools^62^ to filter SNPs and retain only biallelic sites, with no more than 40% missing data, a quality > 30, and with minor allele frequencies > 0.05. The final dataset comprised 100 individuals and 6,700,191 variants (Supp. Table S7). We checked for individual relatedness using *PLINK 1.*9^63^ and found no directly related individual pairs (all genetic relationship values were < 0.2). Finally, we conducted the genome-wide association study (GWAS) on color morph using *PLINK 1.*9^63^ and visualized the result of the GWAS using the R package *qqman*^64^.

### FoxD3 Expression Analysis

RNAseq reads were aligned to the annotated reference genome using HISAT2^65^ and StringTie2^66^. A transcriptome-wide gene count matrix was then created using the script *prepDE.py3* provided on the StringTie website. Subsequent analyses were conducted in the *R* environment^67^. We used the *R* package *edgeR*^68^ to filter and normalized our data prior analysis. We filtered out genes which had a count inferior to 1 count-per-million (cpm) in at least half of the samples, and applied a trimmed mean of M-values (TMM) normalization of the data. We then calculated a contrast matrix and corrected for Poisson count noise using the *makeContrast* and *voom* functions of the R package *limma*^69^, respectively. Developmental stage was introduced in the model as a co-variate. Finally, we compared *FoxD3* expression levels between green and brown individuals (n = 33, one sample per individual; see Supp. Table S9), as well as between sections of dorsal skin within and outside the vertebral stripe within individuals (n = 14 individuals, 2 samples per individual; Supp. Table S9), using the *eBayes* function (Supp. Table S5). We also compared green and brown individuals within, and outside the stripe for wide-striped individuals (n = 14). For tadpoles and metamorphs, which do not display their adult coloration yet, we predicted adult coloration based on their GCP locus genotype.

### Tests for Selection in Ptychadena robeensis

To detect potential signatures of selection acting on the GCP locus, we computed several population genomics statistics on our region of interest as well as across the entire genome using the higher coverage *Ptychadena robeensis* dataset (n=10). We estimated overall and morph-specific nucleotide diversity (π), as well as and inter-morph sequences divergence (DXY) and relative differentiation (FST) using *Pixy*^70^. *Pixy* explicitly accounts for missing genotypes when computing genetic diversity statistics and therefore is more robust to missing data than other methods^70^. As *Pixy* requires invariant sites information, we generated a VCF containing invariant sites using the -*-all-sites true* option in the *GenotypeVCF* function of *gatk*^61^. We filtered out sites with > 40% missing data, and computed statistics over 3 kb sliding window with 1kb overlap between windows.

To search specifically for signatures of long-term balancing selection, we used the summary statistics Tajima’s D^71^, as well as β of *BetaScan2*^72^. Tajima’s D tests for deviation from neutral molecular evolution by comparing the number of segregating sites and nucleotide diversity in a genomic region. Null Tajima’s D values are consistent with neutral evolution, negative values indicate directional selection (positive or negative), and positive values indicate an increase in intermediate frequency polymorphism, consistent with balancing selection. We computed Tajima’s D on a 3 kb sliding window with 1 kb overlap using *vcf-kit*^73^. The β statistic, on the other hand, is useful to detect long or very long-term balancing selection (>25N_ancestral_ generations, where N_ancestral_ is the effective population size of the ancestral population in which the alleles first appeared)^74^. Loci with β values above the 99^th^ percentile of genome-wide values can be considered as likely under long-term balancing selection. We computed β across the entire genome on 3 kb windows in order to determine a threshold based on the top 1% β values and compared the β values in our region of interest to the threshold value.

### Recombination rate mapping in Ptychadena robeensis

Balancing selection can be favored by local suppression or reduction of recombination rate due, for example, to a chromosomal inversion^3,40,42^. To assess whether the maintenance of polymorphism at the GCP locus was favored by local reduction of recombination, we computed the recombination rate map along the genome using *LDhat*^75^ and the higher-coverage *P. robeensis* dataset including ten individuals (8.33X average coverage). Sites with >50% missing data, and non-biallelic sites were filtered out. We ran the *interval* program with a block penalty of 5 for 10,000,000 iterations, sampling every 5,000 iterations. We used a pre-computed lookup table with parameters closest to our dataset (n = 50 haploid individuals and theta per site = 0.001) to generate a new lookup table using *lkgen*. We compared local recombination values at the GCP locus with the chromosome-wide average.

### Comparative genomics

Together with *Ptychadena robeensis*, three species (*P. nana, P. erlangeri,* and *P. levenorum*) form a monophyletic group within the Ethiopian *Ptychadena* radiation that shares the same green/brown polymorphism. We compared the genomic region corresponding to our association peak in *P. robeensis* between green and brown individuals in all three species (*P. nana*: 4 green, 6 brown; *P. erlangeri*: 5 green, 6 brown; *P. levenorum*: 2 green, 3 brown). We found consistent variations between morphs across species, although our low sample size did not allow for a genome-wide association study approach. We found multiple shared polymorphic sites across the four species concentrated in the region of our association peak in *P. robeensis* (Fig. 3F). We computed summary statistics along 3kb sliding windows with a 1kb overlap for all species using *Pixy*^70^ and the same parameters as for *P. robeensis*. Similar to *P. robeensis*, all three species exhibited higher values of π and D_XY_ in the region of interest (Supp. Fig. S4). F_ST_ values increased in the region of interest in all species but *Ptychadena levenorum*, which may be due to the lower sample size (n=5) for this species compared to the other two (n=10).

Because haplotypes were not identical across species, and in order to establish their relatedness, we aimed to reconstruct a haplotype phylogeny. As the locus is intergenic and in order to determine the exact boundaries of the region with shared polymorphism, we conducted a *TWISST*^76^ analysis including all four species and both green and brown individuals (eight groups in total). We phased the genomes of our four species using Beagle^77^ with default parameters. We computed topology weighting on 50 SNPs overlapping sliding windows along chromosome 7. We then summed the weights for all topologies where green haplotypes grouped together across species versus where they did not, and plotted the summed weights. The summed weight of topologies grouping the green individuals together increased significantly in the region matching our association peak in *P. robeensis* (Supp. Fig. S8). We used a 0.05 weight threshold to establish the boundaries of our region of interest, which was 3.38 kb in length. We then combined, genotyped and phased the genomes of the 12 *Ptychadena* species of the radiation using the same parameters for *gatk* and *beagle*, and adding 6 *P. amharensis* individuals, as well as of one individual of *P. beka, P. cooperi, P. delphina, P. doro, P. goweri, P. harenna,* and *P. neumanni* to the previous dataset. Finally, we reconstructed a phylogeny of this 3.38 kb genomic region for the phased haplotypes of our 12 *Ptychadena* species using *RaxML-NG*^78^ and 100 bootstrap iterations. The region counted 762 SNPs after filtering out indels, non-biallelic sites, low quality (Q<30), and extreme depth coverage (DP˂2 and DP>25).

## Data availability

All genetic data used in this article is available at NCBI SRA under accession ###. The reference genome for *Ptychadena robeensis* (assembly GCA_036250615.1) is available at NCBI under accession PRJNA722913.

## Code availability

Scripts used for the analyses of the data are available on GitHub (https://github.com/SandraGoutte/frog_color_polymorphism).

## Supplementary Tables

**Table S1. Color and habitat data used in the comparative analysis.**

**Table S2.** Comparison of fit for four Mk models of evolution for the green coloration in anurans. ER model: transitions between “fixed green”, “polymorphic” and “not green” states occur at the same rate. SYM model: transitions between two states occurring at the same rate regardless of the direction but differ amongst state pairs. ARD model: all transitions occur at different rates. POLY model: transitions from “not green” to “fixed green” and vice-versa must go through the “polymorphic” state. Number of parameters estimated (k), log likelihood, Akaike Information Criterion (AIC), and Akaike weights are given.

| Model | k | log.Lik | AIC | Akaike weight |
| --- | --- | --- | --- | --- |
| ER | 1 | -1179.22 | 2360.44 | 0.00 |
| SYM | 3 | -1121.12 | 2248.23 | 0.00 |
| ARD | 6 | -1008.07 | 2028.14 | 0.54 |
| POLY | 4 | -1010.21 | 2028.43 | 0.46 |

**Table S3.**
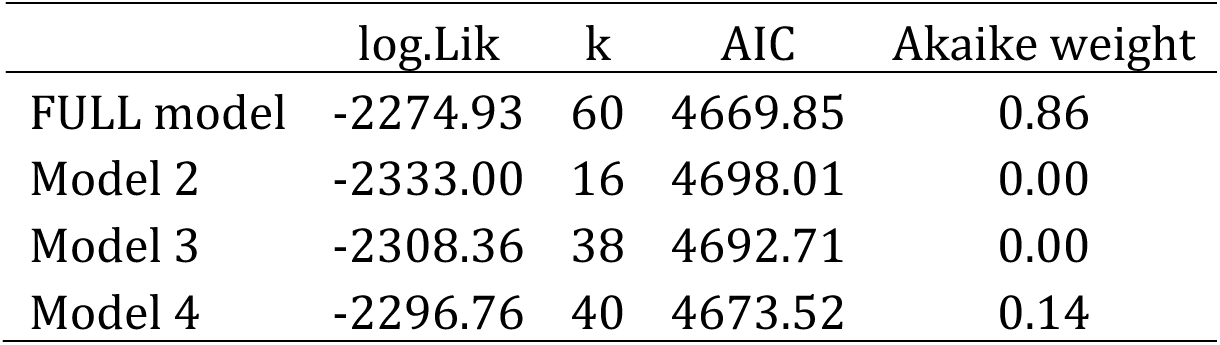
Comparison of fit for four Mk models of green coloration and habitat joint evolution. In order to limit the number of estimated parameters, only the habitat transitions from the Reduced model 2 (see Table S6) are allowed, and fixed color states (“fixed green” and “not green”) must transit through a polymorphic state. Additionally, simultaneous habitat and color transition are not allowed. FULL model: all transition rates are different. Model 2: transition rates between habitats are all different, transitions rates between color morphs are all different, but color state transition rates are the same across habitats. Model 3: all color state transition rates are different within and across habitats, but transitions between habitats have the same rate regardless of the color state. Model 4: all color state transition rates are different but independent of habitat, and habitat transitions rates are all different and vary according to color state.

|  | log.Lik | k | AIC | Akaike weight |
| --- | --- | --- | --- | --- |
| FULL model | -2274.93 | 60 | 4669.85 | 0.86 |
| Model 2 | -2333.00 | 16 | 4698.01 | 0.00 |
| Model 3 | -2308.36 | 38 | 4692.71 | 0.00 |
| Model 4 | -2296.76 | 40 | 4673.52 | 0.14 |

**Table S4.** Comparison of four speciation/extinction (MuSSE) models. NULL model: speciation and extinction are independent of color state, transitions between color states occur at different rates. ER NULL model: speciation and extinction are independent, transitions amongst color states occur at a single rate. ER MuSSE model: speciation and extinction are dependent of color state, transitions amongst color states occur at a single rate. MuSSE model: speciation and extinction are dependent of color state and transitions between color states occur at different rates. Degrees of freedom (Df), Akaike Information Criterion (AIC), difference in AIC compared to the lowest AIC value (Δ AIC), Akaike weight, log likelihood, χ^2^, and P-value of the likelihood ratio test compared to the simplest model (ER NULL) are given.

| | Df | AIC | $\Delta$ AIC | Akaike weight | log.Lik | $\chi^2$ | Pr(>\|$\chi$\|) |
| --- | --- | --- | --- | --- | --- | --- | --- |
| ER NULL | 3 | 39687.25 | 1059.04 | $1.1 \cdot 10^{-230}$ | -19840.62 | | |
| NULL | 6 | 39221.10 | 592.89 | $1.8 \cdot 10^{-129}$ | -19604.55 | 472.15 | $< 2.2 \cdot 10^{-16}$ |
| ER.MuSSE | 7 | 39458.19 | 829.98 | $5.9 \cdot 10^{-181}$ | -19722.10 | 237.05 | $< 2.2 \cdot 10^{-16}$ |
| MuSSE | 10 | 38628.21 | 0.00 | 1 | -19304.11 | 1073.04 | $< 2.2 \cdot 10^{-16}$ |

**Table S5.** Differential expression of *FoxD3* in the dorsal skin of *Ptychadena robeensis*. Models, number of samples (N), average *FoxD3* expression over the samples included in the model (Avg. Expr.), moderated t statistics (t), P-value (P), and log2 fold change (logFC) computed using the functions *voom* and *eBayes* are given for each contrast.

|  | N | Df | Avg. Expr. | t | P | logFC |
| --- | --- | --- | --- | --- | --- | --- |
| *Foxd3* ~ *color* + *dvpt_stage* | 33 | 28 | 2.2 |  |  |  |
| green vs. brown |  |  |  | 2.43 | 0.02 | 1.18 |
| *Foxd3* ~ *colorzone* + *dvpt_stage* + *indiv.* | 28 | 11 | 2.52 |  |  |  |
| brown_IN vs. brown_OUT |  |  |  | 0.21 | 0.83 | 0.07 |
| green_IN vs. green_OUT |  |  |  | 3.03 | < 0.01 | 1.67 |
| *Foxd3* ~ *colorzone* + *dvpt_stage* | 28 | 21 | 2.52 |  |  |  |
| green_OUT vs. brown_OUT |  |  |  | 0.4 | 0.69 | 0.2 |
| green_IN vs. brown_IN |  |  |  | 3.5 | < 0.01 | 1.72 |

**Table S6.** Comparison of fit for six Mk models of habitat evolution. ER model: transitions amongst all six habitat categories occur at a single rate. SYM model: transitions between two habitats occur at the same rate regardless of the direction, but differ amongst state pairs. ARD model: all transitions amongst habitats are allowed and occur at different rates. Ordered model: only transitions between adjacent habitats are allowed: arboreal ˂-˃ arboreal-terrestrial ˂-˃ terrestrial ˂-˃ terrestrial-aquatic ˂-˃ aquatic and terrestrial ˂-˃ fossorial-terrestrial. All allowed transitions occur at different rates. Reduced models include all transitions with a rate ≥ 0.001 (Reduced model 1) or > 0.001 (Reduced model 2) in the ARD model (see Supplementary Fig. S5).

|  | k | log.Lik | AIC | Akaike weight |
| --- | --- | --- | --- | --- |
| ER | 1 | -1617.44 | 3236.88 | 0.00 |
| SYM | 15 | -1460.17 | 2950.34 | 0.00 |
| Ordered | 10 | -1504.70 | 3029.39 | 0.00 |
| Reduced 1 | 14 | -1439.29 | 2906.59 | 0.30 |
| Reduced 2 | 12 | -1440.47 | 2904.93 | 0.69 |
| ARD | 30 | -1428.00 | 2915.99 | 0.00 |

**Table S7. *Ptychadena robeensis* samples used for the Genome-wide association study**

**Table S8. Samples used for the comparative genomics analysis**

**Table S9. *Ptychadena robeensis* samples used for the RNAseq analysis.**

## Supplementary Figures

**Figure S1.**
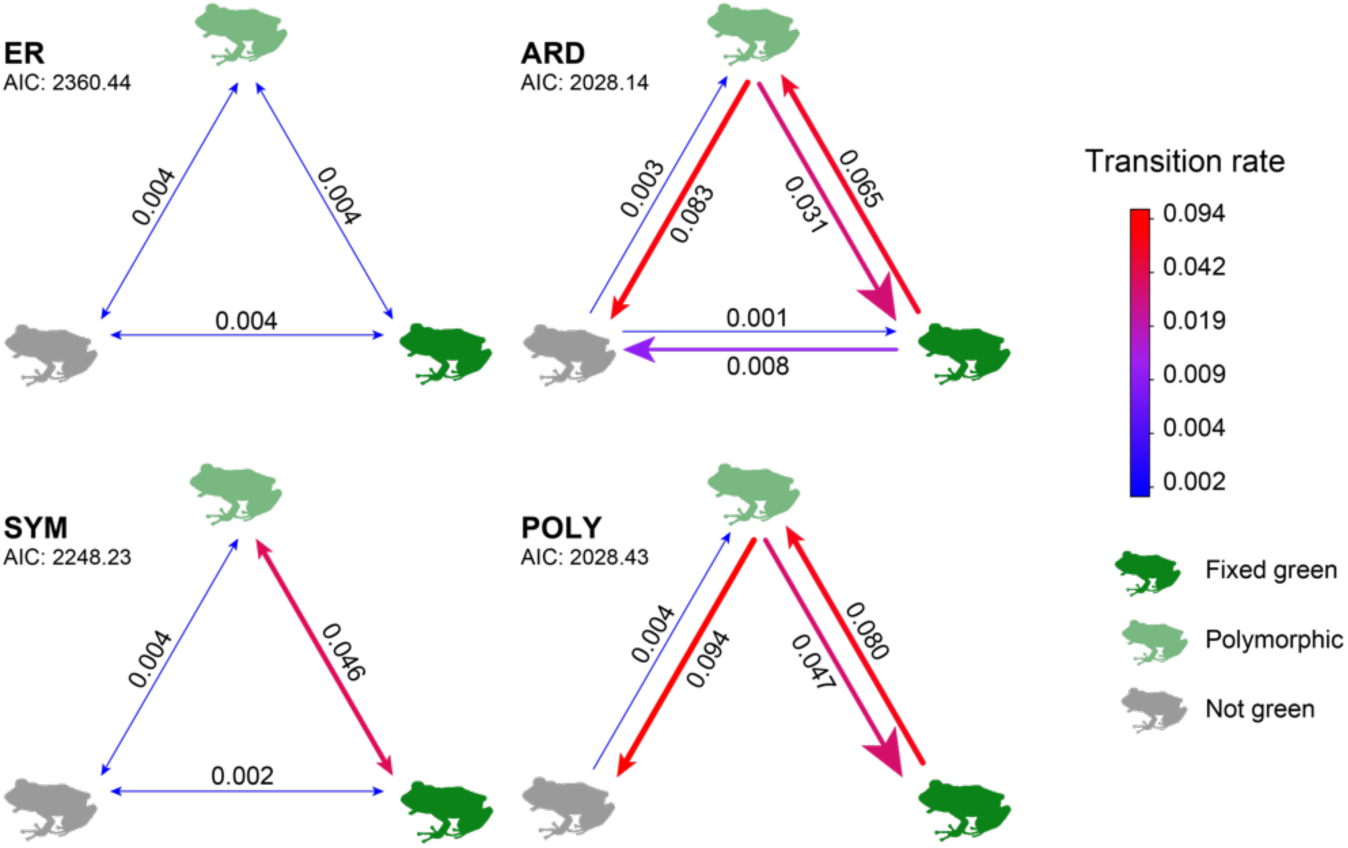
Comparison of four Mk models of evolution for the green coloration in anurans. The ARD (all rates different) and POLY (color changes transit through the “polymorphic” state) models better fit the data than the ER (equal rates) and SYM (symmetrical rates) models. Goodness of fit does not significantly differ between the ARD and the POLY models (Likelihood ratio = 4.2935 (df=2), P = 0.1169).

**Figure S2.**
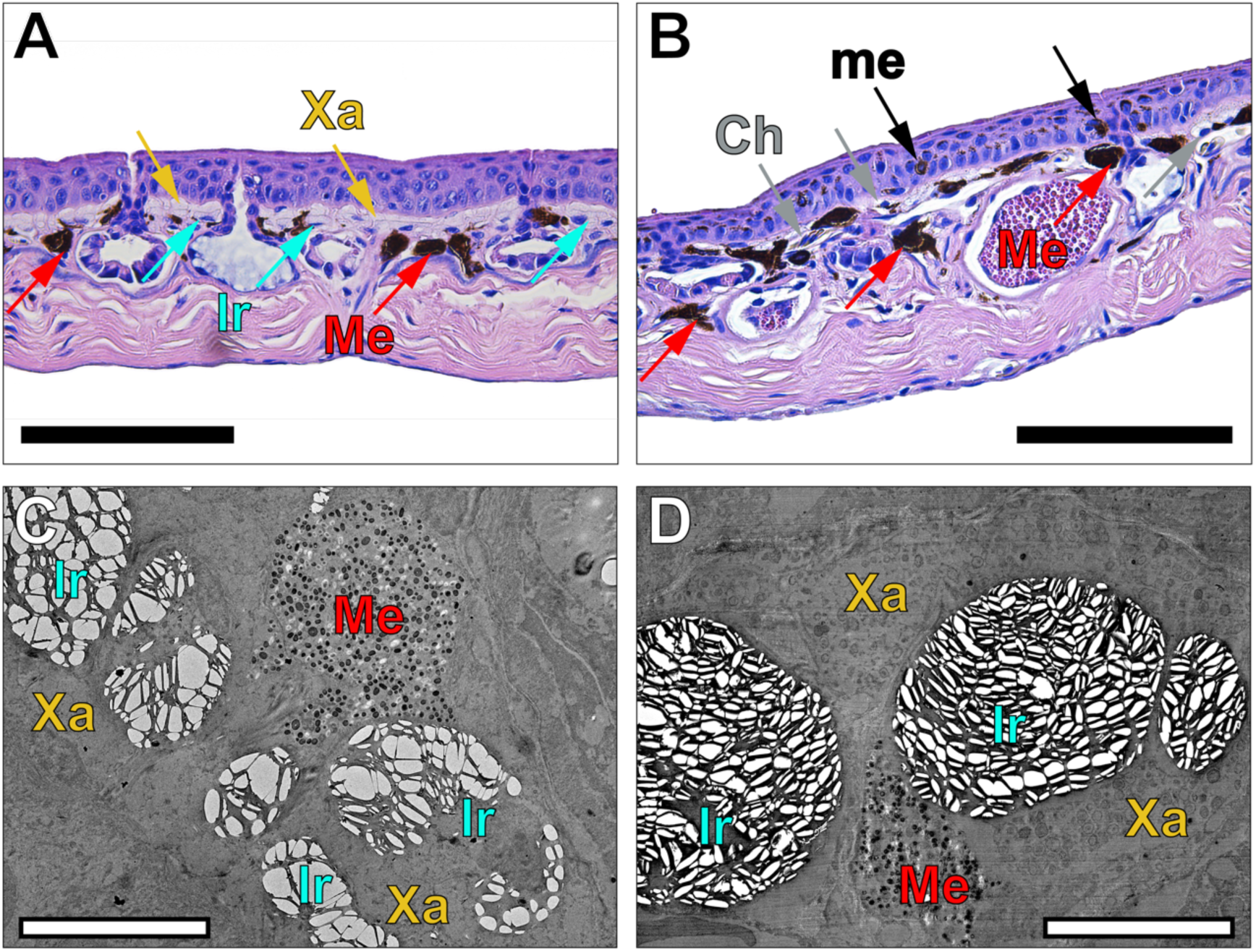
Cellular and sub-cellular structures of green and brown dorsal skin in *Ptychadena robeensis*. **A &B** Histological sections of green (**A**) and brown (**B**) dorsal skin. Scale: 100 µm. Green skin contains two continuous and regular layers of light chromatophores (xanthophores (Xa) and iridophores (Ir)) and few melanophores (Me), while brown skin harbors a greater density of melanophores, fewer and less organized light chromatophores (Ch), and epidermal melanosomes (me). **C & D** Electron micrographs of green (C) and brown (D) skin. Scale: 10 nm. All three types of chromatophores are present in both green and brown skin, but the size and shape of iridophores’ guanine crystals differ between the two morphs. Pigments within the xanthophores also appear to be darker in brown skin. See supplementary text.

**Figure S3.**
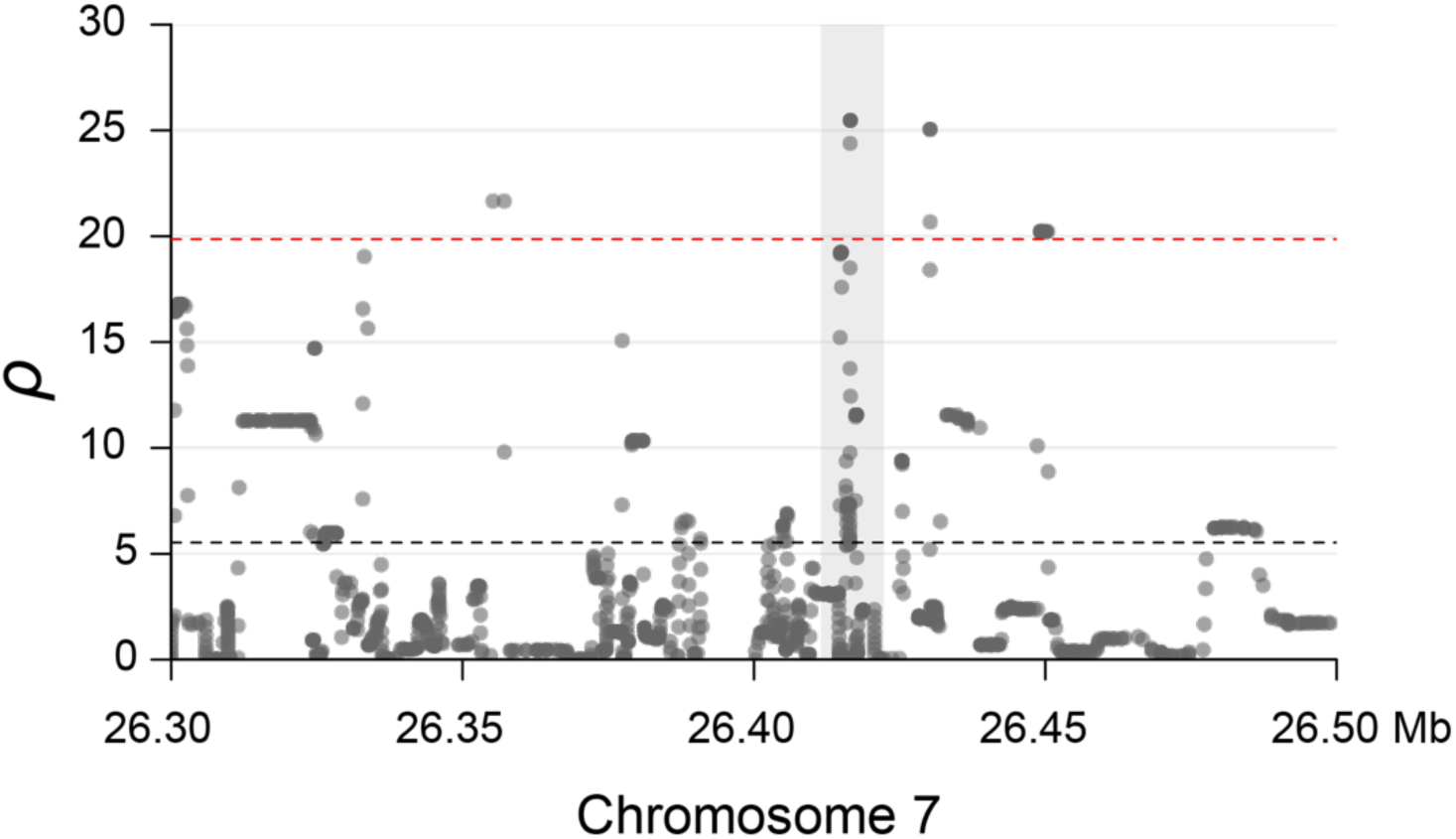
Recombination rate (*ρ*) in the GCP region of *Ptychadena robeensis*. Population-wide recombination map computed along the genomes using the ten individuals high-coverage (8.33X average coverage) dataset and LDhat^75^. Grey-shaded area corresponds to the GCP-associated region (chromosome 7, position: 26,411,500- 26,422,300). Horizontal dashed lines represent the average (black) and 90^th^ percentile (red) *ρ* values.

**Figure S4.**
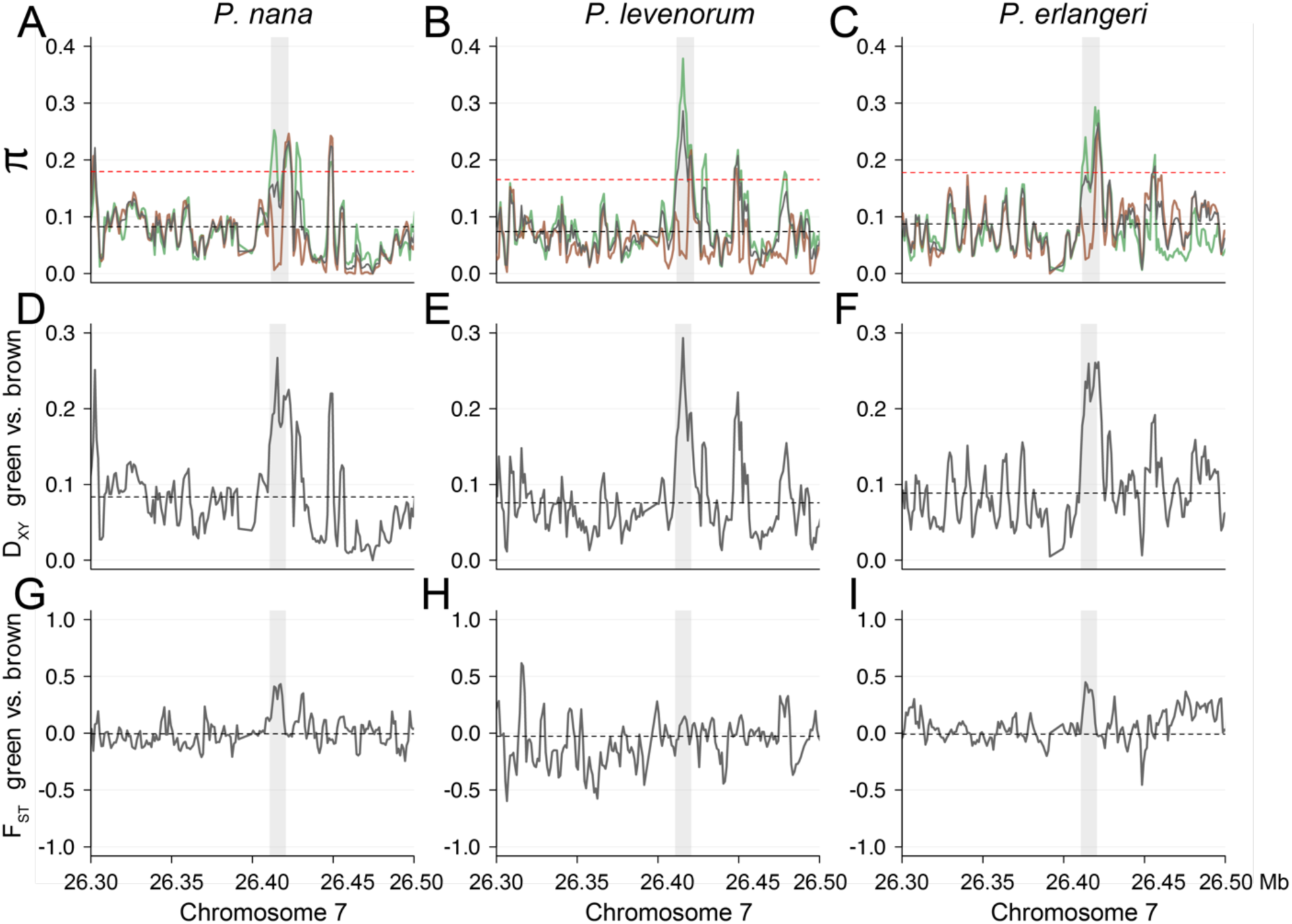
Signatures of balancing selection in the GCP region of three polymorphic *Ptychadena* species. All summary statistics were computed on 3 kb sliding windows with 1 kb overlap using *Pixy* ^70^(n=10 *P. nana*, n=5 *P. levenorum*, and n=11 *P. erlangeri*; 6.33X average coverage). Grey-shaded area corresponds to the GCP-associated region (chromosome 7, position: 26,411,500-26,422,300). Black horizontal dashed lines represent the genome-wide average statistics’ value. **A-C** Population-wide (grey), brown individuals (brown), and green individuals (green) nucleotide diversity (π). Red horizontal dashed line represents the genome-wide 95^th^ percentile of nucleotide diversity. **D-F** Absolute genetic divergence (D_XY_) between green and brown individuals. **G-I** Relative differentiation (F_ST_) between green and brown individuals.

**Figure S5.**
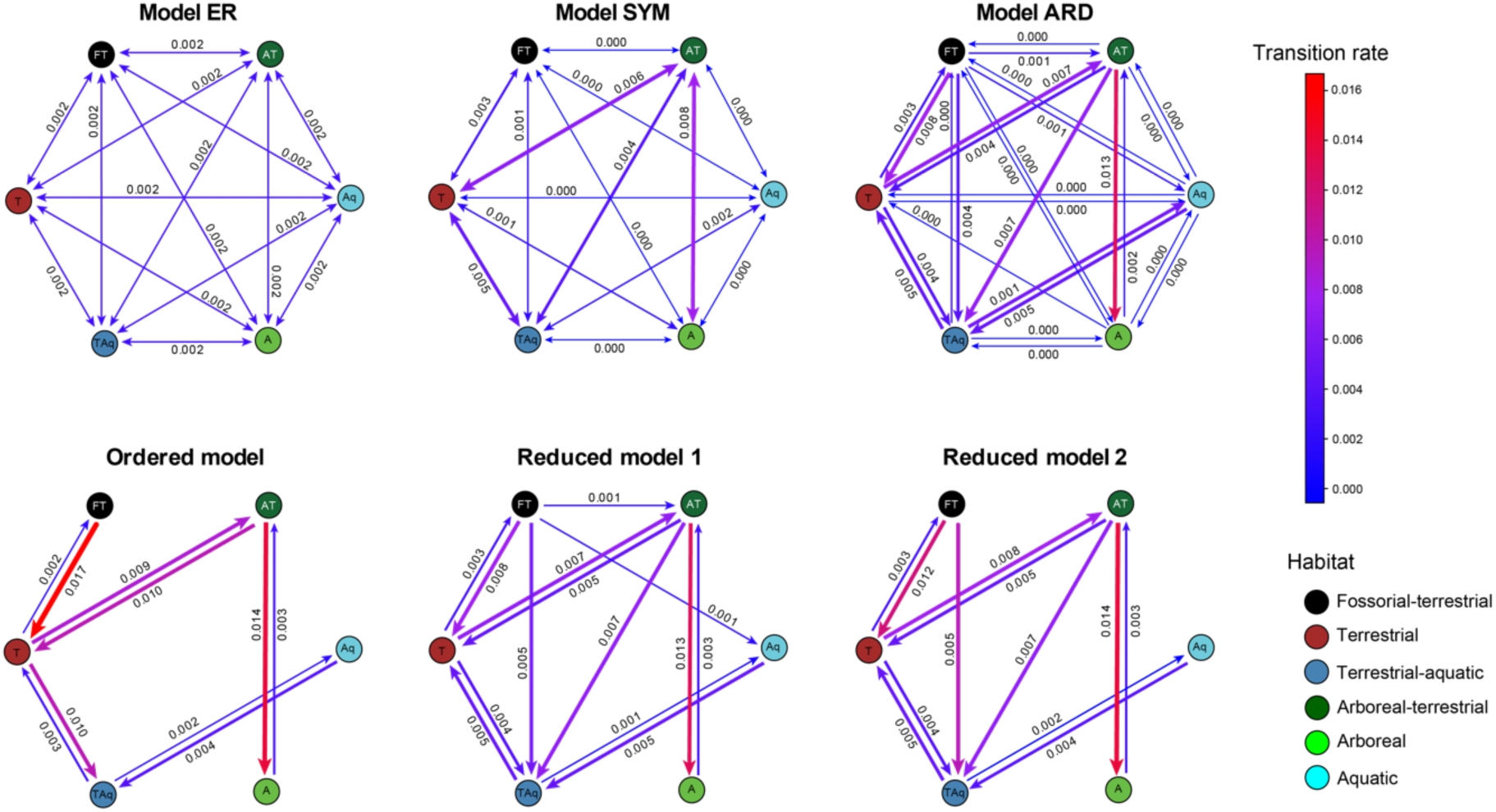
Six Mk models for habitat evolution. Model ER estimates a single transition rate for all possible habitat transitions (k = 1). Model SYM estimates one transition per habitat pair, independently of the direction of the transition (k = 15). Model ARD estimates all possible transition rates (k = 30). The ordered model considers only transitions between adjacent habitats (e.g. aquatic ˂-˃ terrestrial-aquatic ˂-˃ terrestrial), and estimates each possible transition rate (k = 10). The reduced models only estimate transitions rates which are ≥ 0.001 (reduced model 1; k = 14), and > 0.001 (reduced model 2; k = 12) in the model ARD.

**Figure S6.**
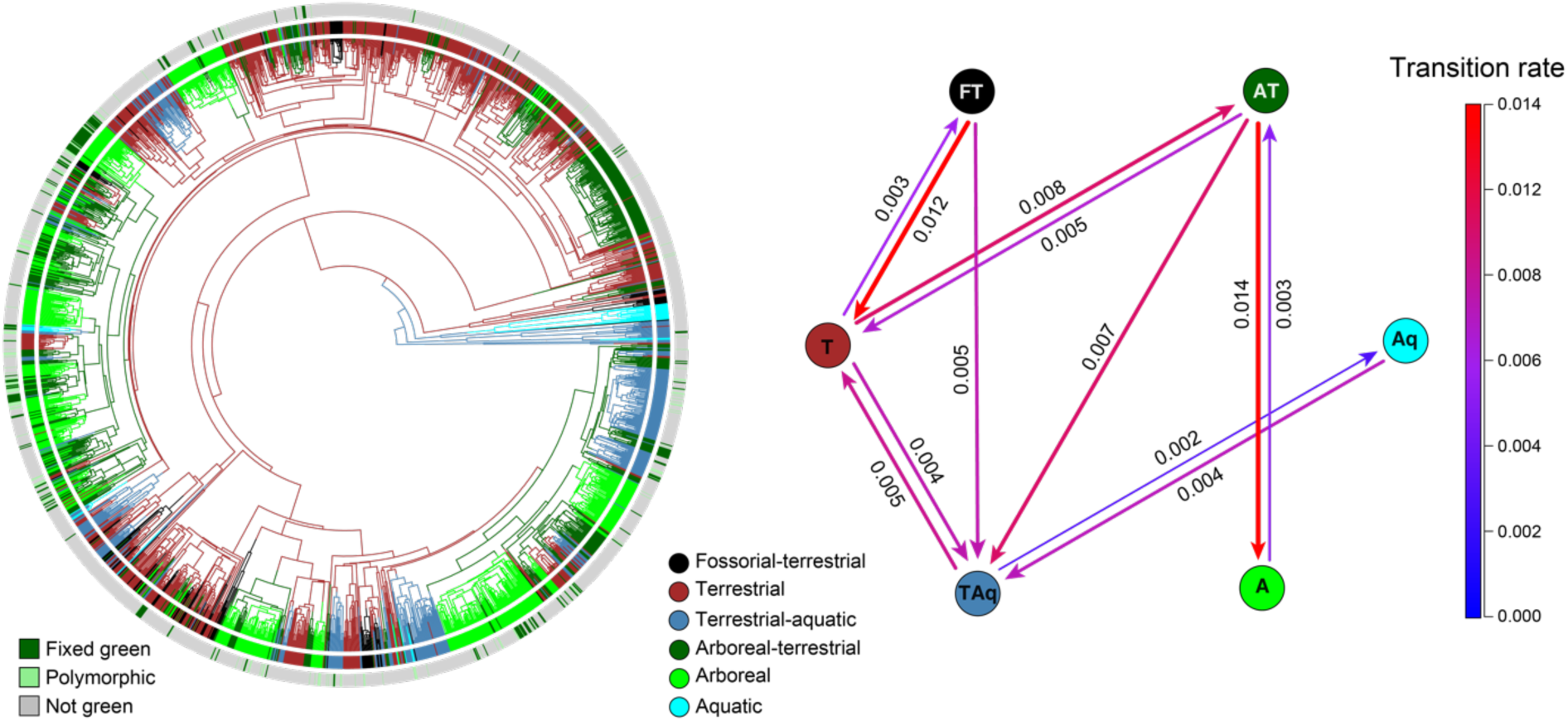
Evolution of habitat preferences in anurans. Left: phylogeny including 1,989 species (phylogeny from^28^) with mapped habitat and green coloration data. Tree branches are colored according to the consensus habitat state established using 1,000 stochastic maps. Right: best-fitting Mk model for transitions amongst the six habitats.

**Figure S7.**
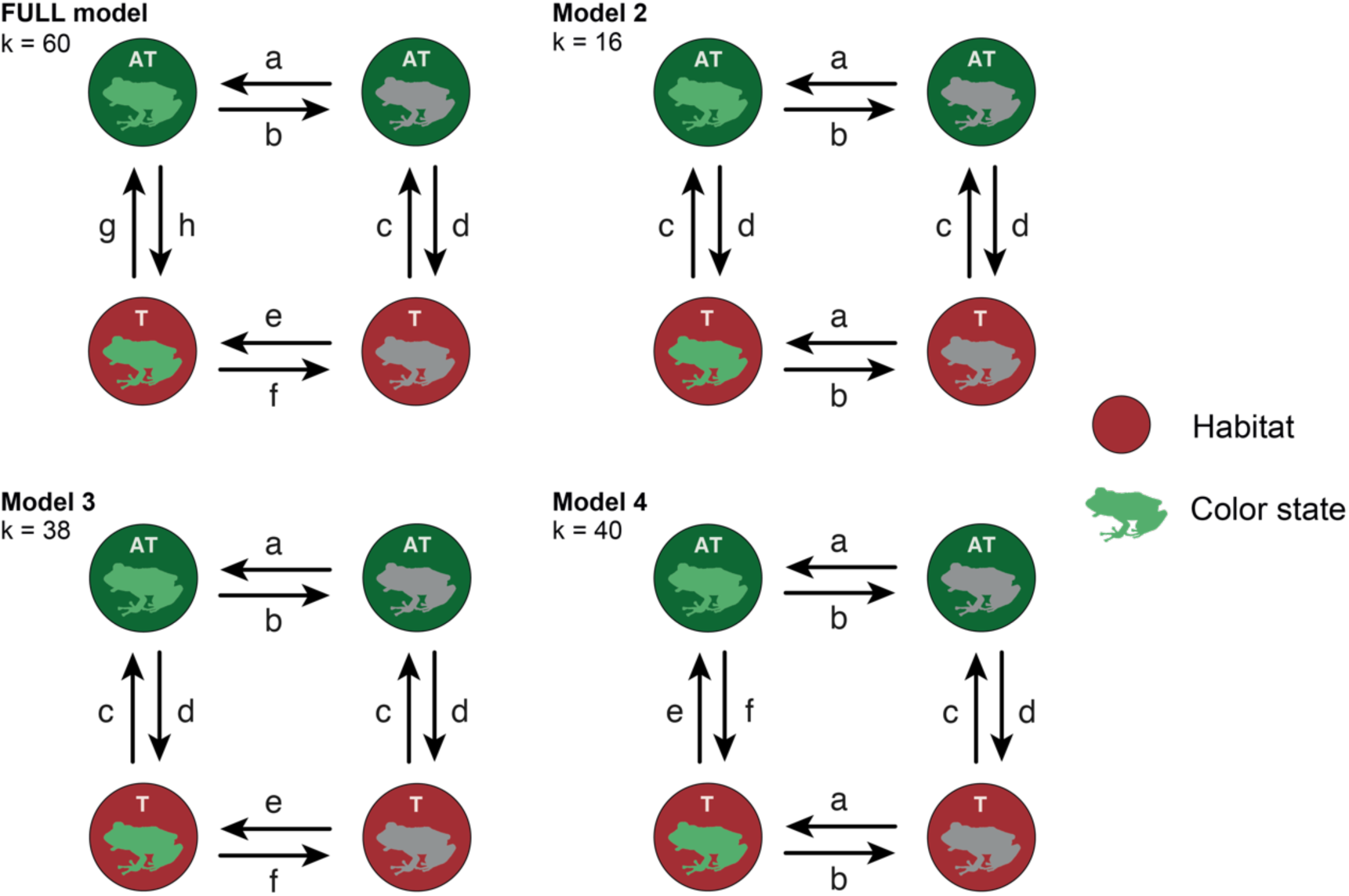
Four models of green coloration and habitat joint evolution. The FULL model considers all transition between color states to occur at different rates and to be dependent on habitat, and all habitat transition rates to be different and dependent on color state (k = 60). Model 2 considers transitions between color states to be independent of habitat and transitions between habitats to be independent of color state (k = 16). Model 3 considers all transitions between color states to be different and dependent on habitat, but habitat transitions to be independent of color state (k = 38). Model 4 considers transitions between color states to be independent of habitat but habitat transitions to be dependent on color state (k = 40).

**Figure S8.**
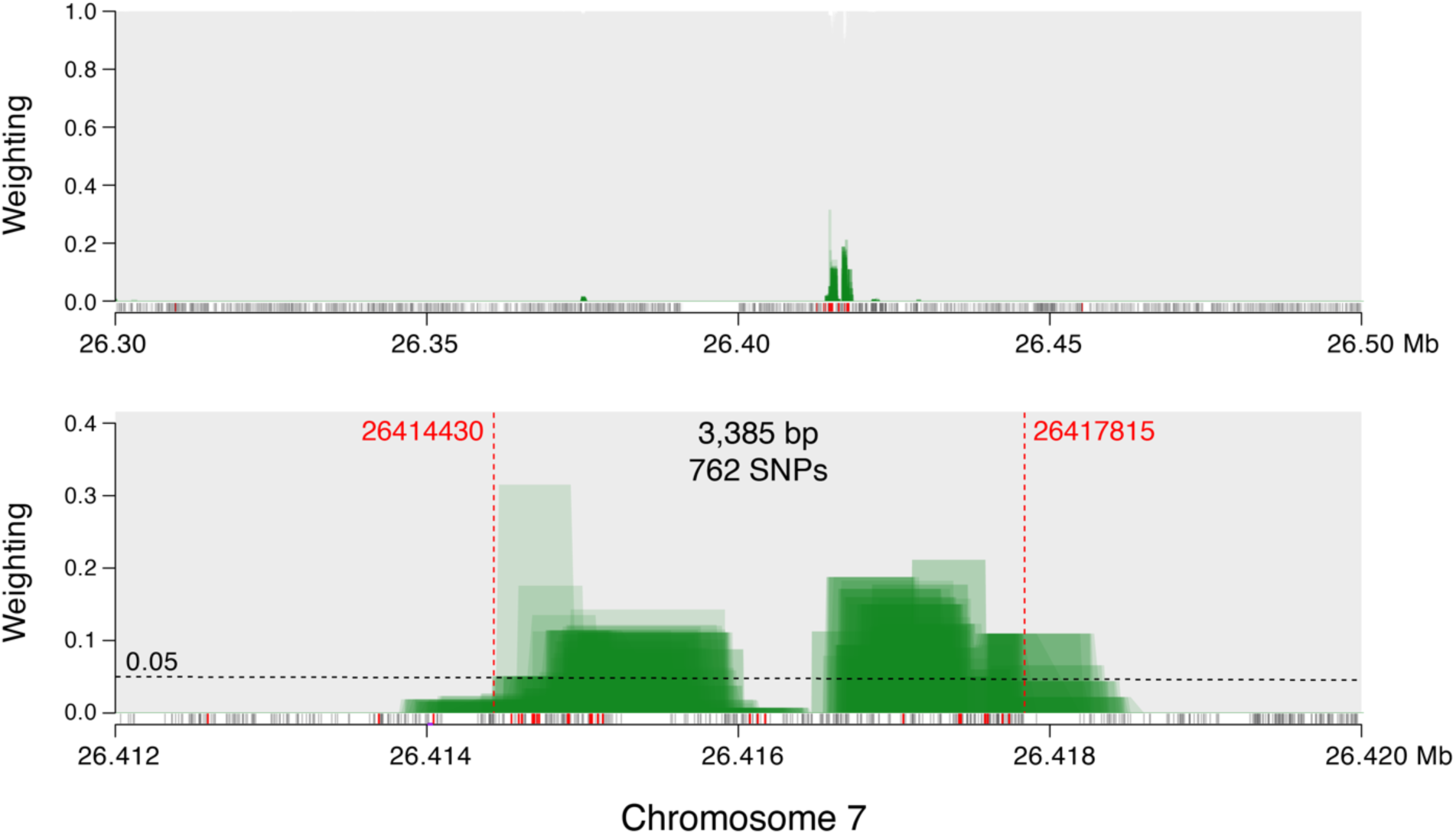
Topology weighting in four polymorphic *Ptychadena* species. Green and brown individuals of four polymorphic *Ptychadena* species (*P. robeensis, P. nana, P. levenorum,* and *P. erlangeri*) were included in the TWISST analysis resulting in eight groups (see text). Weights of topologies grouping the green individuals separately from the brown ones were summed and are displayed in green. All other topologies are displayed in grey. Solid grey bars at the base of each panel indicate SNPs. Solid red bars indicate SNPs with shared polymorphism across the four species. The window used for the haplotype phylogeny reconstruction (Fig. 4) is indicated by vertical dashed lines in the bottom panel.

## Supplementary text

### Histology and transmission electron microscopy of Ptychadena robeensis dorsal skin

We examined the cellular and subcellular structures of green and brown dorsal skin in *Ptychadena robeensis* using light and transmission electron microscopy. Dorsal skin sections were extracted from seven preserved adult specimens (three green and four brown), two green juveniles and three tadpoles *Ptychadena robeensis*. For light microscopy, the skin samples were embedded in paraffin blocks and sections of 4 µm thickness were produced. The sections were stained with hematoxylin-eosin (HE) and chromatophores were examined using a Leica DMI 6000 B microscope. We used the following staining protocol: xylene 2 × 2 min, absolute ethanol 2 × 2 min, 95% ethanol 1 min, 70% ethanol 1 min, water for 2 × 30 s, hematoxylin 5 min, water for 2 × 30 s, bluing reagents 30 s, water 30 s, 95% ethanol 30 s, Eosin 3 min, 95% ethanol 30 s, absolute ethanol 2 × 30 s, xylene for 2 × 30 s. For transmission electron microscopy (TEM), sections within and outside the stripe of four adult individuals (two green and two brown) were extracted and processed separately. Skin samples were stained in 1% osmium tetroxide in 0.1 M PB buffer for four hours, dehydrated through a series of graded ethanol baths from 50% to 100% for 15 min each, then two baths of 100% acetone for 15 min each. Samples were then placed in 50:50 mixture of acetone and embedding resin (Embed 812, Electron Microscopy Sciences) overnight followed by a bath of 100% resin for four hours. Samples were finally embedded in fresh 100% resin and placed in a 60° C oven for 16 hours for polymerization. Blocks were sectioned with glass knifes in 100 nm thickness sections and placed on 200 mesh copper grids for imaging in TEM or scanning transmission electron microscopy (STEM) microscopes.

Examination of histological sections revealed that green skin had two continuous layers of light-colored chromatophores, and few melanophore in contracted state underneath (Supp. Fig. S2A). Brown skin had light chromatophores interspersed with melanophores, which could be in contracted or dispersed states (Supp. Fig. S2B). In brown skin, the density of melanophores was greater than in green skin, and epidermal melanosomes were common, while they were absent in green skin (Supp. Fig. S2B). Transmission electron microscopy revealed that all three types of chromatophores (xanthophores, iridophores and melanophores) were present in brown and green skin, but iridophores formed a continuous layer and the guanine crystals contained within the iridophores were larger and rounder in green skin compared to brown skin, which may explain their difference in reflectance^35^ (Supp. Fig. S2C & S2D). Additionally, pigments within the xanthophores appeared lighter in green skin compared to brown skin (Supp. Fig. S2C & S2D). Both cellular and sub-cellular structures thus appear to play a role in the difference in color in the dorsal skin of *Ptychadena robeensis*.

## Notes

### Competing Interest Statement

The authors have declared no competing interest.

